# Quantifying the number of translatable transcripts through the use of OMICs involved in post-transcriptional regulation

**DOI:** 10.1101/2022.06.20.496876

**Authors:** Juan Ochoteco Asensio, Jos Kleinjans, Florian Caiment

## Abstract

Transcriptomics is nowadays frequently used as an analytical tool to study the extent of cell expression changes between two phenotypes or between different conditions. However, an important portion of the significant changes observed in transcriptomics at the gene level is usually not consistently detected at the protein level by proteomics. This poor correlation between the measured transcriptome and proteome is probably mainly due to post-transcriptional regulation, among which miRNA and circRNA have been proposed to play an important role. Therefore, since both miRNA and circRNA are also quantified by transcriptomics, we proposed to build a model taking those factors into account to estimate, for each transcript, the fraction of transcripts that would be available for translation. Using a dataset of cells exposed to diverse compounds, we evaluated how our model was able to improve the correlation between the assessed transcriptome and proteome expression level. The results show that the model improved the correlation for a subset of genes, probably due to the regulation of different miRNAs across the genome.

## Introduction

The central dogma of molecular biology states a straightforward flow of information: for a gene to transmit the information it contains, its DNA needs to be transcribed to RNA to produce the desired proteins. Each of these biological pools of molecules has its field of study for both their characterization and quantification, which are commonly known as omics. Transcriptomics, for example, refers to the large-scale study of transcripts, and the same applies to proteomics (proteins) or metabolomics (metabolites).

Proteins, being the functional molecules of the cell, are of high interest to be analyzed, as they accurately represent the phenotypic changes of the studied cell or tissue. Unfortunately, the technology’s sensitivity and reproducibility behind proteomics, namely mass spectrometry, is currently still limited, especially when the aim is to get a wide protein expression analysis. This is mainly due to the mass spectrometer only being able to measure a fraction of the eluted peptides, hence giving as an output a slight portion of the entire population of proteins^1–3^. To circumvent this issue, an alternative strategy often used is the expression analysis of their precursors: the transcripts. Transcriptomics is indeed often performed as the surrogate to analyze different disease states or cell conditions. Even so, the fact that transcripts are indeed the cause and origin of proteins is not reflected by a high correlation between both omics^4–8^. This is rather unsurprising, due to the multiplicity and complexity of the factors playing a role in post-transcriptional regulation, in addition to intrinsic variable characteristics of both molecules (such as half-life^9^), and some of those regulatory factors are even transcripts themselves.

One of such factors is microRNAs (miRNAs), whose effect on the expression levels of proteins, although mild, has been characterized for more than a decade^10,11^. MiRNAs are short non-coding RNA sequences playing a role in translation regulation^12^. They contain the so-called “seed” region^13^, a short sequence with perfect Watson-Crick complementarity with their target (primarily to their 3’ UTR region). The binding between the two molecules impairs the translation of the target^14^. Due to the short length of the seed region, a single miRNA often targets several transcripts, and a single transcript is targeted by numerous miRNAs. This inhibitory relationship may differ depending on the level of expression of the miRNA and its target molecules, the number of regions in a single mRNA complementary to the miRNA present, and the combinatorial effect of several miRNAs targeting the same transcript. Taking into account these parameters is necessary to have a better comprehension of the mechanisms behind post-transcriptional regulation.

Another element to be considered in post-transcription regulation is circular RNAs (circRNAs). circRNAs are transcripts that possess a circular structure due to the covalent binding of their 5’ and 3’ ends^15,16^, hence obtaining a circular structure that bestows them resistance to exonuclease activity^17^. One of their recently discovered functions is described as ‘miRNA sponge’^18^. This refers to their ability to contain several copies of short sequences that are complementary to the miRNAs’ seed region. Consequently, they also play a role in the regulation of translation machinery by competing against other transcripts for miRNA binding^19^. The more miRNAs are captured by circRNAs, the more transcripts will be available for translation.

CircRNAs are not the only targets competing for miRNA, as other (long) non-coding RNAs (ncRNAs) can also present a seed target for some particular miRNAs^20^. This collection of both coding and non-coding transcripts can be conceptualized as a single group of targets for a shared miRNA, which led them to be generally called competing endogenous RNAs (ceRNAs)^21^.

Thus, in the current age of sequencing, connecting more precisely transcriptomics to the actual phenotype of the cell is a major challenge. Currently, many publications base their biological interpretation on the differentially expressed genes in two groups of samples as transcriptomics is more sensitive and allows the assessment of almost all possible RNA molecules in a single experiment. In this context, we wondered if it would be possible to integrate all available transcriptomics information into a model able to estimate the level of translation of any expressed coding transcript, namely, the fraction of translatable transcripts (TrT). This value, computed for each protein-coding transcript, is an estimation of the number of transcripts that are free to be translated after taking into account the aforementioned post-transcriptional regulation.

To design and assess the model, we used an *in vitro* dataset obtained from human cardiac microtissues exposed to a range of compounds at different doses, which were analyzed with proteomics, RNA-Seq (ribo-depleted libraries), and miRNA-Seq methods. From the RNA sequencing data, we first identified and quantified different transcript biotypes: protein-coding, non-coding, miRNA, and circRNAs. The proposed model was then applied to generate the TrT score from all possible interactions between those molecules. In this manuscript, we analyze the possible benefits of such an approach compared to the state-of-the-art methods for gene expression analysis.

## Methods

### Samples

The analyzed data consists of 3D microtissues containing stem-cell-derived cardiomyocytes and fibroblasts in a 4:1 ratio obtained from InSphero. These microtissues were exposed to 8 compounds: Fluorouracil (5FU), Amiodarone (AMI), Celecoxib (CEL), Docetaxel (DOC), Doxorubicin (DOX), Epirubicin (EPI), Mitoxantrone (MXT), and Paclitaxel (PTX), in addition to a control group (Untreated/UNTR). The dosing profile was established via the use of the physiologically based pharmacokinetic (PBPK) modeling software PK-Sim, to simulate exposure levels under physiological conditions^22^. For each compound, 2 doses were applied: Therapeutic and Toxic. The exposures were done in triplicates, and the data extraction was performed at 7 time-points: 2, 8, 24, 72, 168, 240, and 336 hours; resulting in 21 data-points per dose. Four samples were excluded due to low sequencing depth: three samples for having too low miRNA sequencing depth (CEL/MXT/PTX_Tox_002_3; toxic dose, time-point 2h, 3^rd^ triplicate) and one for its RNA-Seq low read count (UNTR_002_3).

### RNA Sequencing (RNA-Seq)

Total RNA from the exposed microtissues was isolated using the Qiagen AllPrep Universal Kit (Cat #80224). Ribo-depletion was achieved by using the Illumina RiboZero Gold kit (Cat #MRZG12324), and the libraries were prepared using the Lexogen SENSE total RNA kit (Cat #009.96). All libraries were then sequenced on an Illumina HiSeq 2000 at 100 bp paired-end at an average coverage range between 20 and 30 million reads. The adaptors were removed through Trimmomatic version 0.33^23^. We used the following parameters: paired-end, ILLUMINACLIP: TruSeq3-PE.fa:2:30:10, LEADING:3, TRAILING:3, SLIDINGWINDOW:4:15, MINLEN:36, HEADCROP:12.

### Proteomics

Proteins were isolated and diluted to a concentration below 0.2M. The peptides were digested by using trypsin, and peptides were cleaned-up using Sep-Pak tC18 cartridges (Waters) according to the manufacturer’s instructions. A vacuum centrifuge was used to dry the peptides, before measuring them on an Orbitrap Fusion mass spectrometer (Thermo Fisher Scientific), which was coupled to a NanoLC-2D HPLC system (Eksigent). To process the raw MS data, Genedata Expressionist software (v.11.0) was used. The noise in the LC-MS peaks was reduced and normalized, and afterward, their properties were obtained (m/z and RT boundaries, m/z and RT center values, intensity). To annotate the individual MS/MS spectra, Mascot 2.6 was used. To group peak clusters, protein interference was used (based on peptide and protein annotations), and, using the Hi3 method, protein intensities were computed.

### miRNA analysis

Starting from the same total RNA isolated for the ribo-depleted libraries, an aliquot was size selected and ligated using the TruSeq Small RNA Library Prep Kit (Illumina®). After sequencing on the HiSeq 2000 at 3.6 million reads per sample (after quantification), we quantified the resulting data using miRge2 (last change: 05/06/2018)^24^. miRge2 used the MirBase database as the reference library (miRBase v22), bowtie-1.1.1^25^ as the mapper, miRge2/sp as the miRge library, human as the species selected, and illumina for the adapter to be removed. The output results were in gff format. Isomirs were not considered for this analysis.

### (non-)coding RNA analysis

The genome version used for all the transcriptomics analyses was the Genome Reference Consortium Human Build 38 (GRCh38.p12). For the identification of circRNAs, we performed a *de novo* prediction from our dataset. For this, we concatenated all RNA-Seq forward data (R1) across all 240 Cardiac samples into a single FASTQ file. Afterward, this file was inputted to two recent circRNA prediction software: circExplorer2^26^ and CIRI2^27^. For circExplorer2, we first used BWA version 0.7.17^28^ to index the genome and align (minimum score to output: 19) the reads to the indexed genome. Afterward, we used circExplorer2 to parse the aligned reads and annotate the circRNAs. For CIRI2, we also required the aligning step via BWA, and then the annotation step through its internal algorithm (-S (single-end), -U 3 -B 13, to set mapping quality thresholds of a junction read and help control False Discovery Rate (FDR)). We then extracted the overlap of identified molecules between both outputs for decreasing the amount of false-positive predictions. The circRNA IDs were comprised of the chromosomal and strand locus of the predicted molecules. Based on this information, we extracted their genetic sequence using BEDTools^29^, obtaining a final circRNA transcriptome library.

We downloaded the transcriptomes for both coding (all cDNA) and non-coding (all ncRNA) transcripts from Ensembl (release 96)^30^. We combined these 2 libraries with our predicted circRNAs one into a single library, which we set as the global transcriptome reference for Salmon^31^, with which we quantified the RNA-Seq data. Salmon output contained both the number of reads and TPM (Transcript per Million) values for each transcript, the latter of which was used for the analysis. The benefit of using TPM values, instead of raw counts, relies on the inherent normalization performed for both sequencing depth and transcript length. If raw reads would have been used instead, the variability across samples and compounds would have made it impossible to optimize the model consistently. Moreover, since our model computes interactions between molecules of different sizes, using read counts would have greatly biased the output.

### CircRNA predictors selection

We selected CIRI2 and circExplorer2 out of 4 possible predictors, which also included find_circ^18^ and circRNA_finder^32^. The parameters for find_circ were set as default, and the samtools version was 1.3. For circRNA_finder, it involved 2 steps: running STAR (-c -- runThreadN 4 --chimSegmentMin 20 --chimScoreMin 1 --alignIntronMax 500000 -- outFilterMismatchNmax 4 --alignTranscriptsPerReadNmax 100000 --twopassMode Basic -- outSAMtype BAM SortedByCoordinate --chimOutType SeparateSAMold -- outFilterMultimapNmax 2) and the post-processing script (--minLen 100). We formatted all the output equally, so that we were able to compare them across tools.

### Interaction tables

To evaluate all possible bindings between the molecules, a table containing all possible interactions between miRNAs and any putative competing ceRNAs was generated, each row representing a unique interaction.For miRNA, both identifiers (IDs) and sequences were sourced from miRBase^33^. For all transcripts (both coding and non-coding), we extracted the Ensembl transcript IDs through biomaRt^34^. We obtained the sequences corresponding to these IDs through SAMtools^35^, which were searched in the genome FASTA file by using the chromosomal coordinates of the transcripts. For mRNAs, we used only the 3’ UTR region.To establish the microRNA target interactions (or MTIs), we used miRanda (version 3.3a)^36^ with the -strict parameter to force strict 5’ seed pairing. We set the score threshold to 140.0 (corresponding to a full 7 base pair seed with no mismatch). We also used –noenergy, disabling the thermodynamics performance.

### Translatable Transcripts (TrT) formulation

We designed a formula (Formula 1) that aimed to estimate the fraction of translatable transcripts based on a basic principle: the number of translatable transcripts is equal to the difference between the total expression of a given protein-coding transcript and its inhibited molecules. The number of inhibited transcripts depends on the number of miRNAs and the probability that these will bind to the target transcript. The probability of a target being targeted by miRNA depends on how abundant the target is in proportion to the number of all the possible targets. For all data analyses, we used R^37,38^ as the main programming language.

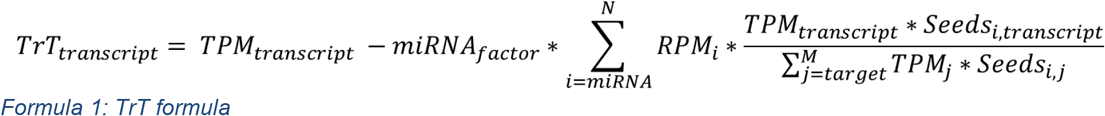

TrT = Translatable Transcripts, TPM = Transcripts Per Million, RPM = Reads Per Million, Seeds = the number of seeds present in a specific transcript targeted by a specific miRNA, N = number of miRNAs targeting the transcript, M = number of targets for a specific miRNA_i_, miRNA_factor_ is a scaling factor, set to 0.1 (TrT model design).

### TrT example

For example, consider a protein-coding transcript expressed at 100 TPM targeted by two miRNAs, expressed at 132.5 and 227.5 Reads Per Million (RPM) respectively. Each of them has a single seed or sequence region presented by the target that they can bind to. The first miRNA can also interact with a single circRNA, expressed at 2 TPM on 3 perfect seed regions, while the second miRNA can bind to an ncRNA with a TPM value of 30 (Equation b).

#### Box 1

TrT example

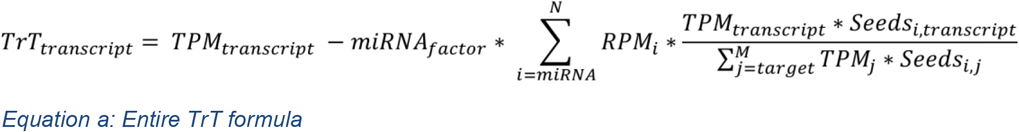

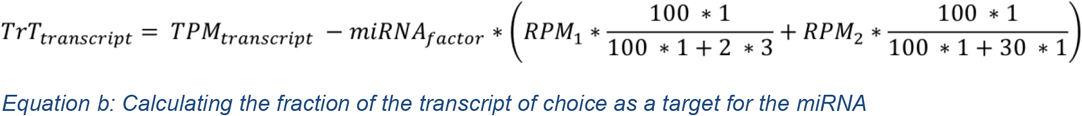

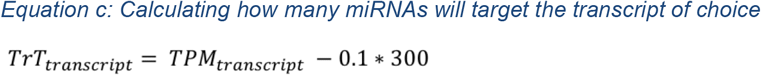

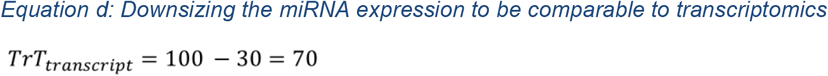

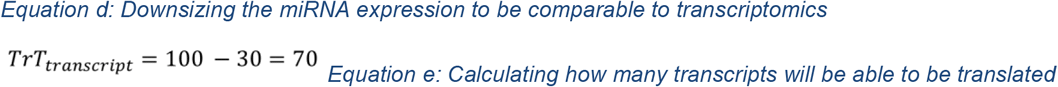

Those miRNAs can bind to any of the aforementioned ceRNAs (other mRNA targets, circRNAs, or ncRNAs). To know how many miRNAs will bind to each of them, we need to know the probability of each interaction. In our study, we hypothesized that the probability that a miRNA will bind to a molecule is directly proportional to how abundant that molecule is in relation to all possible targets (0.94 and 0.77 in this example, Equation c). Consequently, we estimate that 94% (28.3) of these miRNA molecules will bind to the target. This number is multiplied by the miRNA factor (Equation d), and we subtract it from the original TPM value of our target, which will be the expected TrT value (Equation e).

### Correlation between Transcriptomics and Proteomics

To perform correlation analysis between our transcriptomics and proteomics dataset, we used the untreated samples of the first 4 time-points (2, 8, 24, and 72 hours) for transcriptomics, and all the normalized (Proteomics) time-points for proteomics (2, 8, 24, 72, 168, 240, and 336 hours).

To be able to correlate protein and transcript expressions, we merged protein-coding transcripts by their protein product, and we grouped all values per time-point while maintaining the same order to perform a Pearson correlation at the time-point level. For 168h, we omitted the first triplicate due to a significant batch effect, correlating only the other 2 triplicates with the other 2 triplicates of all other time-points. We did the same with the RNA-Seq sample of UNTR_002_3, because of the low sequencing depth, as mentioned before. The graphical displays were generated thanks to the ‘corrplot’ package^39^.

### Differential Expression Analysis

#### Proteomics

To obtain differentially expressed proteins (DEPs), we followed the steps performed by Selevsek et al (2020). Briefly, we first log-2 transformed all the proteomic expression values. We then calculated the median for each control sample. Afterward, we calculated the median of all the medians (MoM) and shifted the control samples so that they all shared this median. For every treatment/dose combination, we determined the set of proteins common in both control and treatment samples. This set of proteins was used to determine the MoM on the normalized control samples. We then shifted the distribution of the data by matching their median values to this MoM. Finally, we performed paired t-tests to evaluate the significance between doses, considering as significant (or DEPs) the ones for which the p-values were lower than 0.05.

#### Transcriptomics

As we aim at evaluating the TrT fraction of the coding transcripts, Salmon mapping output was filtered out for non-coding transcripts. All remaining coding transcript expression values were summed based on their gene of origin. While usually, differential expression analyses are performed via a dedicated sequencing analysis pipeline on raw read count using a pipeline specific for negative binomial distribution (such as DESeq2 or EdgeR), this approach could not be applied here. Indeed, in our case, the values to be compared (TPM and TrT) did not allow such pipelines, either because the input was expected to be un-normalized^40^ and/or because other normalization steps were performed instead^41^. We then log-2 transformed both TPM and TrT values, which, like proteomics, also presented originally a negative binomial distribution. We performed t-tests between therapeutic and toxic doses. Due to the high percentage of significant observations and multiple testing, we also applied an FDR/BH p-adjustment. We identified a differentially expressed gene (or DEG) as a gene with a p-adjusted value lower than 0.05

### Biological Interpretation with GOrilla

Using as input the lists of genes ranked by increasing p-adjusted value (for each comparison and compound), we ran a gene ontology enrichment analysis with GOrilla^42^ using the ‘Single ranked list of genes’ mode. We studied the GO terms of interest and analyzed some of their genes’ expression (both in TPM and TrT). We also explored how the MTIs (using the miRTarBase^43^ list) related to such genes (with at least weak evidence of having a regulatory effect) were expressed in comparison to what TPM and TrT were representing.

### Sensitivity, Specificity, and Accuracy

A prediction is normally evaluated by contrasting it to the reality it is trying to model. Such evaluation focuses on different aspects of the predictor: how many of the positives that have been predicted are real positives (sensitivity), how many of the predicted negatives are indeed negative (specificity); and how many cases are correct out of all cases, independently of whether they are positive or negative (accuracy), making use of binary classification terms. In our case, proteomics was considered the true condition, while transcriptomics was the predicted condition. True or false referred then to the correct or incorrect representation of proteomics by transcriptomics, while positive or negative referred to the presence or absence of differential expression, respectively. For example, if both proteomics and transcriptomics showed differential expression in the same manner, such gene was considered a true positive. If transcriptomics did not present a differential expression while proteomics did, it was considered a false negative. Once these terms were defined, the calculation of such evaluators was performed according to the standard statistical measures of a binary classification test performance.

## Results

### Test dataset for TrT model assessment

To develop a TrT model, we used a dataset as input derived from a 3D cardiomyocyte culture, from which both transcriptomics (ribo-depleted and small RNA libraries) and proteomics (LC-MS) data were generated. The cell cultures were composed of 8 individual compound treatments, in addition to an untreated control. For every compound, 3 replicates were measured in 7 time-points (2, 8, 24, 72, 168, 240, and 336 hours) for every dose (therapeutic and toxic), resulting in 42 samples per compound. This dataset, of a total of 240 samples, presented the added value to have been generated by the same technician, and all proteomics and transcriptomics samples were generated from the same run, thus reducing an important source of bias. The three different technologies (RNA-Seq, miRNA-seq, and proteomics) were quantified based on state-of-the-art procedures (Methods). While our RNA-Seq data analysis quantified the protein-coding and non-coding transcripts available in the Ensembl database, circular RNA identification required additional steps in addition to *de novo* prediction.

### CircRNA predictors selection

CircRNAs prediction software are known for generating a high level of false positives^44^. To identify the circular RNAs in this transcriptomics dataset, we thus decided to use an overlapping approach between several tools. For this, we considered the following four tools: CIRI2^27^, find_circ^18^, circRNA_finder^32^, and CIRCexplorer2^26^. When determining the optimal overlap, we interpreted the amount of uniquely predicted identities by a tool as an indicator of the false-positive ratio. We could observe that most of the circRNAs were only predicted by circRNA_finder, which suggested this algorithm has the highest rate of false positives. The other 3 tools showed similar false-positive ratios, so to compare them we focused on the overlap that maximized the number of identified molecules: circExplorer2 and CIRI2. The list of predicted circRNA molecules was then added to the reference transcriptome used to quantify all expressed RNA molecules from the ribo-depleted libraries.

#### General Expression Analysis

Such transcriptome was structured in the following manner: 126831 of the identifiers were non-coding RNA (57.37%), 85108 were protein-coding RNAs (38.50%), and 9138 were circRNAs (4.13%) (Figure 2A).

**Figure 1:**
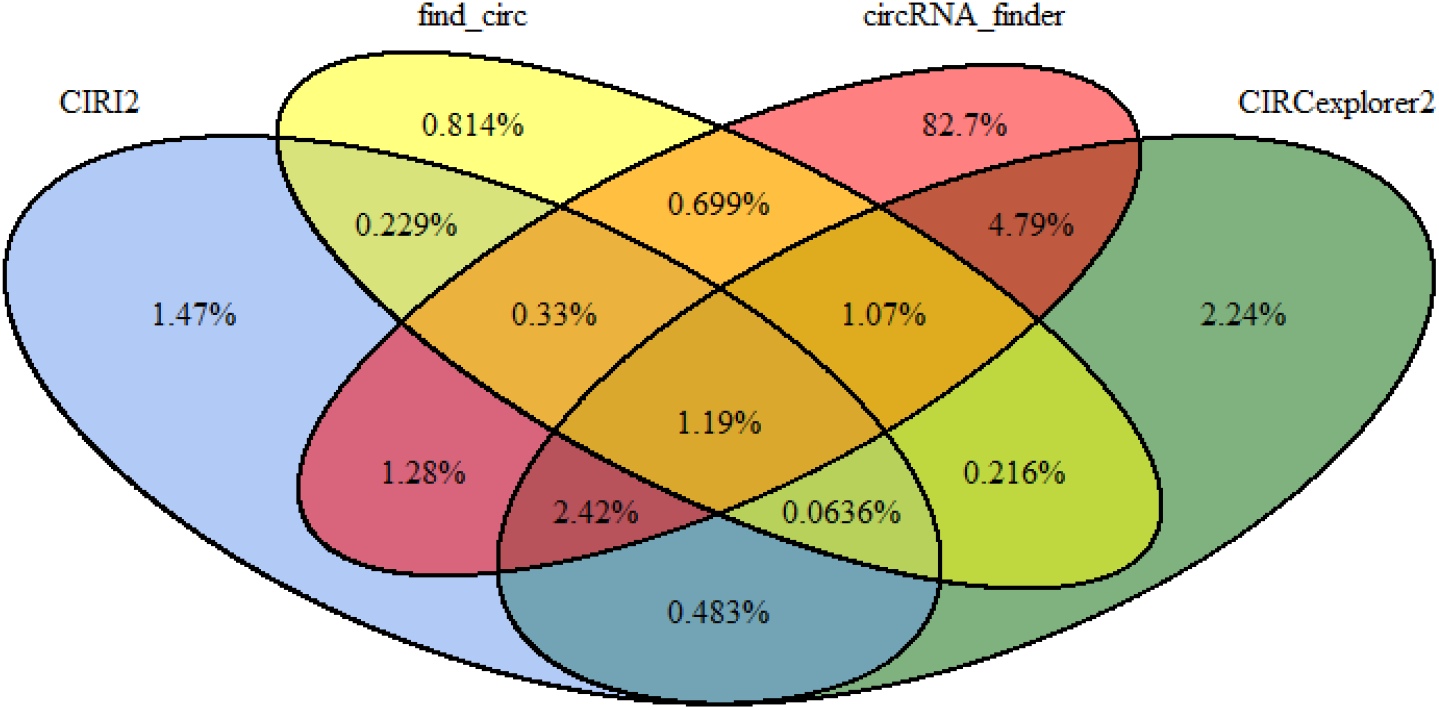
Venn Diagram between 4 circRNA prediction tools output.

**Figure 2:**
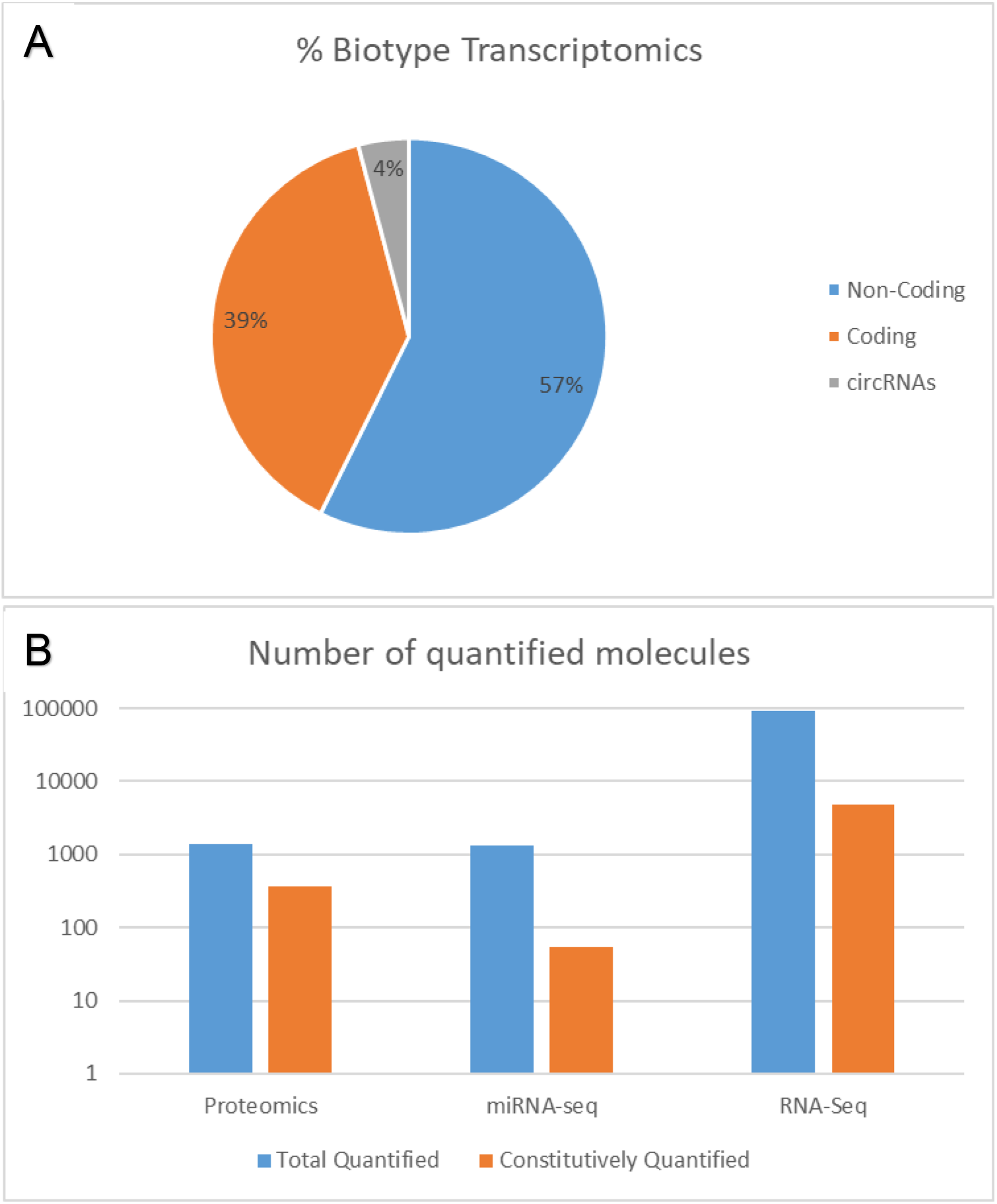
Quantification results across different omics. **A:** Percentage of each biotype present in the Transcriptomics library.**B:** Number of quantified molecules in the Control condition. Total Quantified refers to the number of molecules that have been at least once quantified by such technology. Constitutively Quantified refers to the number of molecules that have been quantified at all samples by such technology.

In control conditions, for RNA-Seq, we detected 92969 transcripts expressed in at least 1 sample > 0 reads, with 4815 (5%) of them present in all 240 samples. In microRNA sequencing (miRNA-seq), out of the 1348 quantified molecules in any sample, only 55 miRNAs (4%) were detected constitutively. In proteomics, there were 1392 proteins quantified, of which 362 (26%) were detected in all samples. As expected, proteomics, having one order of magnitude less quantified molecules (both in Total and Constitutively) than RNA-Seq (**Figure 2**B), was the limiting factor when comparing both. Also, when comparing proteomics and miRNA-seq, although they have similar total quantification numbers, the difference to their constitutively quantified shows that the former has a higher proportion of highly expressed molecules, while miRNA-seq displays a broader representation of all molecules (in addition to its more volatile/temporary function).

### TrT model design

With all individual RNA molecules from the transcriptome characterized and quantified, we investigated the possibility to develop a model to predict the expected translatable fraction (TrT) of any given coding transcript. For this, we hypothesized that taking into account as many post-transcriptional regulation factors would be fundamental. Therefore, we conceptualized a model based on several key features of the various biological factors available. First, miRNAs decreased a transcript’s chance of being translated. An increase in miRNA expression (controlling for all other conditions) would then decrease the number of mRNAs available for translation^45^. Second, the probability that a miRNA transcript interaction (MTI) occurred is directly proportional to the expression level of a target in proportion to the expression level of all possible targets. Lastly, both coding and non-coding RNAs (including circRNAs) that were able to interact with a miRNA inhibiting the coding transcript of interest could work as ceRNA: the higher the expression of a single target, the lower the inhibition for the rest of the targets^46–48^. In other words, we assumed that a given miRNA would be equally distributed among all its possible targets (counting in the number of individual seeds), and only the fraction available to interact with our gene of interest should be considered to have an inhibiting effect.

The central concept behind this model was that the number of transcripts to be translated was the difference between the total amount of transcripts (in TPM) and the ones that would not be translated due to miRNA inhibition. The number of transcripts that would be inhibited was then based on two main factors: the level of expression of each miRNA that had the mRNA as a target (RPMi), and the probability that each miRNA would interact with the target. We hypothesized that the probability that a miRNA would bind to a target was proportional to how prevalent that target was compared to all possible targets. A target did not always equal a single molecule (a circRNA could present several seed regions/targets). For this reason, we represented a target by its total amount of seed regions for a specific MTI (transcript expression level times its number of seed regions).

Because miRNA libraries present a smaller density than ribo-depleted RNA libraries, the raw number of reads generated for the miRNAs were in general higher than the levels of the rest of transcripts. This difference did not allow us to subtract one to another directly, and miRNA expression should be scaled to a comparable level of the coding RNA transcripts. To know which value should be given for this scaling factor, we first analyzed which factors would be optimal for TrT, that is, in which miRNA had enough power to show a difference between TrT and TPM, but not so powerful that it reduced all expression values to zero. We searched such optimal value by evaluating a range of values between 0 and 1 (0 leading to no miRNA effect at all, and 1 leading to no scaling), and selected the values in which TrT maximized the correction benefit in comparison to TPM (taking proteomics as a reference). The optimal range we observed was between 0.1 and 0.27. Besides, initial investigation of a newly developed sequencing method named Combo-Seq, which allows sequencing both transcripts and miRNAs altogether in a single library preparation, showed us that the proportion observed of miRNA represented around 10% of all transcripts sequenced. Consequently, we decided to set the miRNA factor to ‘0.1’ (Translatable Transcripts (TrT) formulation)

### Correlation between Transcriptomics (TPM/TrT) and Proteomics

Having our method described and finished, we applied our method to all the aforementioned data. Afterward, we aimed to compare it with the state of the art TPM to investigate in which manner TrT could be an improvement. Initially, we wanted to confirm the initial issue (transcriptomics does not accurately represent the proteome), and compare such results with TrT. For that purpose, using the untreated samples, we performed a correlation analysis between proteomics and TPM, followed by its analogous analysis between proteomics and TrT.

The correlation analysis confirmed the low correlation between Transcriptomics (TPM) and proteomics values (Figure 3A), the average correlation of which was 0.39 (± 0.06 SD). There was no specific time interval between both omics that presented an improved correlation. For example, 24h proteomics and transcriptomics showed the highest correlations, but at the same time, 2h proteomics and transcriptomics presented one of the lowest. Even so, the correlation values were not randomly distributed, but substantially influenced by which proteomics time-point they were related to. As an example, time-point 24h presented the highest values regardless of the transcriptomics time-point (avg.: 0.46), while time-point 2 hours presented the lowest (avg.: 0.26). To a lower extent, a similar effect was observed for the transcriptomics time-points. When evaluating TrT at the same level, we saw a similar low correlation with proteomics (Figure 3B), confirming the fact that a representative portion of transcripts (80%) presented equal values between TPM and TrT, due to not presenting any predicted inhibition by any miRNA expressed in our dataset.

**Figure 3:**
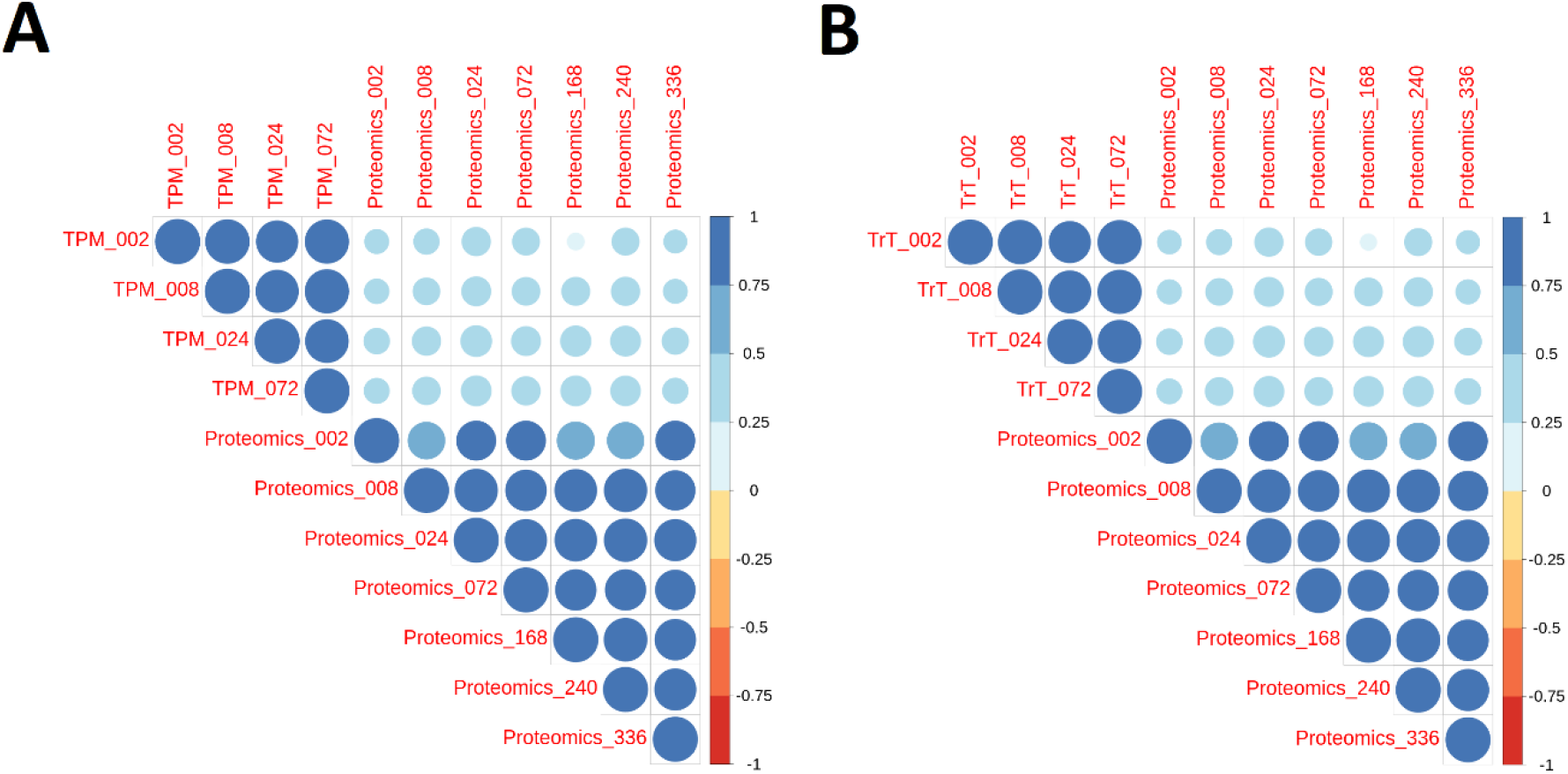
Correlation analysis between Transcriptomics and proteomics. **A:** TPM vs Proteomics. **B:** TrT vs Proteomics. TPM: Transcripts Per Million. TrT: Translatable Transcripts. Timepoint samples are compared across each other (from 2 to 336 hours). The darker and bigger a circle is, the higher the absolute value of correlation it represents. Blue stands for positive correlation, red for negative correlation.

### Differential Expression Analysis

The high similarity between both correlation analyses was not surprising, as we predicted and formulated for miRNA to have generally a small regulatory effect. Even so, we hypothesized that for a portion of the transcripts regulated by miRNA, TrT would be a better proxy for proteomics. To verify this, we ran our formula through all our data. Later, we investigated in which cases traditional transcriptomics was falsely reporting as a proxy by taking proteomics as a reference. The analysis involved the identification of differentially expressed molecules across doses. Two possible scenarios unfolded: there could be a change in transcriptomics not reflected in proteomics, and vice versa: a change in proteomics that was not reflected in transcriptomics. For example, in fluorouracil’s (5-FU) exposure, 5.48% of genes presented a change in transcriptomics and not in proteomics, while 4.02% of them did not present a change while proteomics did. Therefore, we focused on the genes for which TrT could correct such cases.

The myosin heavy chain 9 (MYH9) gene was one of them. We could observe a significant increase (p.value < 0.05) (Figure 4A) of the protein at a Toxic dose of 5-FU, while for TPM it was not (p.value > 0.05) (Figure 4B). TrT, on the contrary, did reflect a significant increase (p. adjusted < 0.05) (Figure 4C).

**Figure 4:**
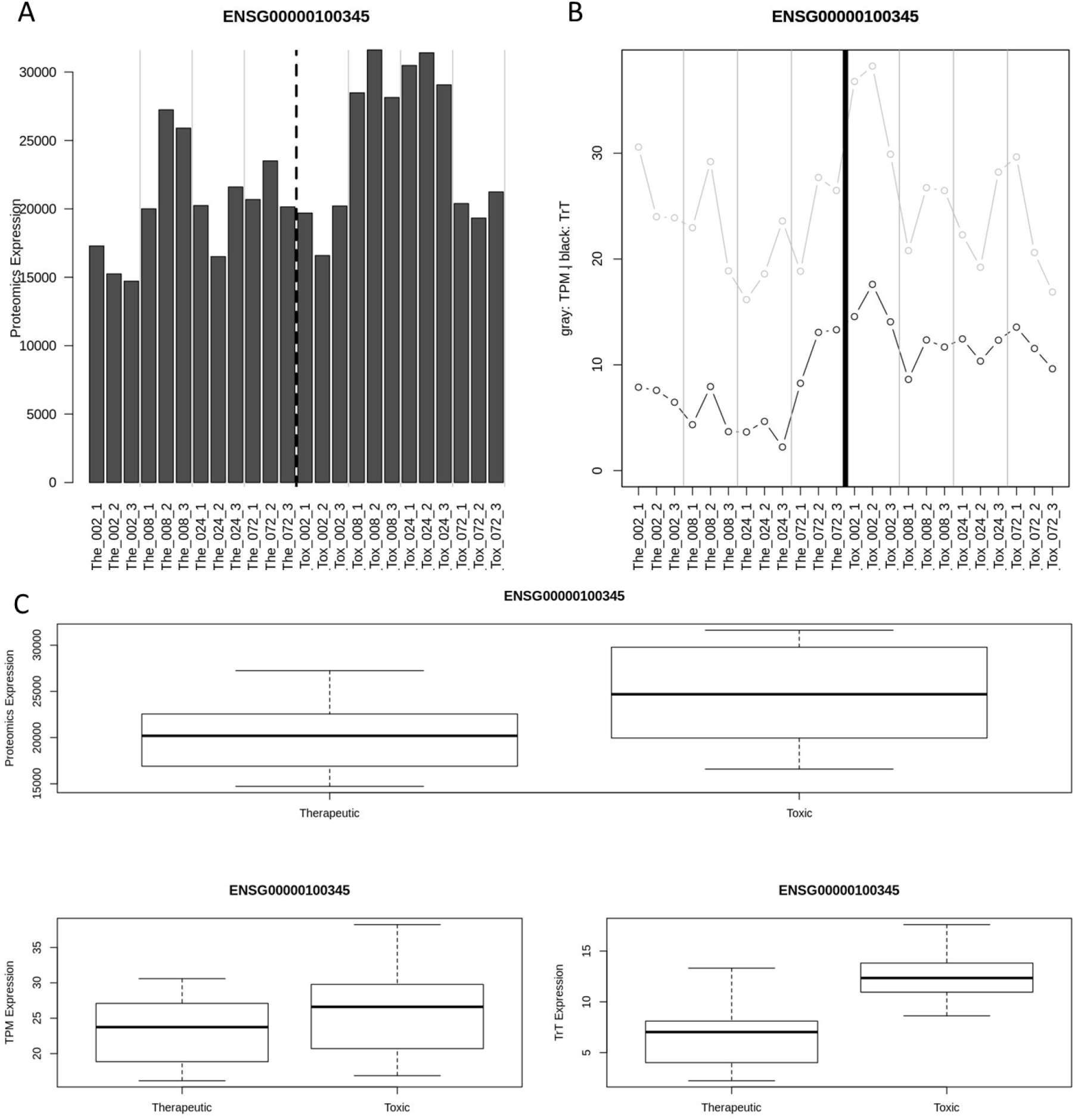
MYH9 expression values for therapeutic and toxic doses of 5-FU. **A:** Proteomics expression values. **B:** TPM and TrT expression values: Gray: TPM, Black: TrT. **C**: Boxplots of the expression values for proteomics, TPM, and TrT. TPM: Transcripts Per Million. TrT: Translatable Transcripts.

We could also find examples of the opposite circumstance. Such was the case for TGFBI (transforming growth factor beta-induced). In this case, proteomics did not show any significant change across doses of Docetaxel (Figure 5A), but TPM was significantly increased (p.adjusted < 0.05) in the toxic dose (Figure 5B). TrT, though, was consistent with proteomics, not showing any significant difference between doses (p.value > 0.05) (Figure 5C).

**Figure 5:**
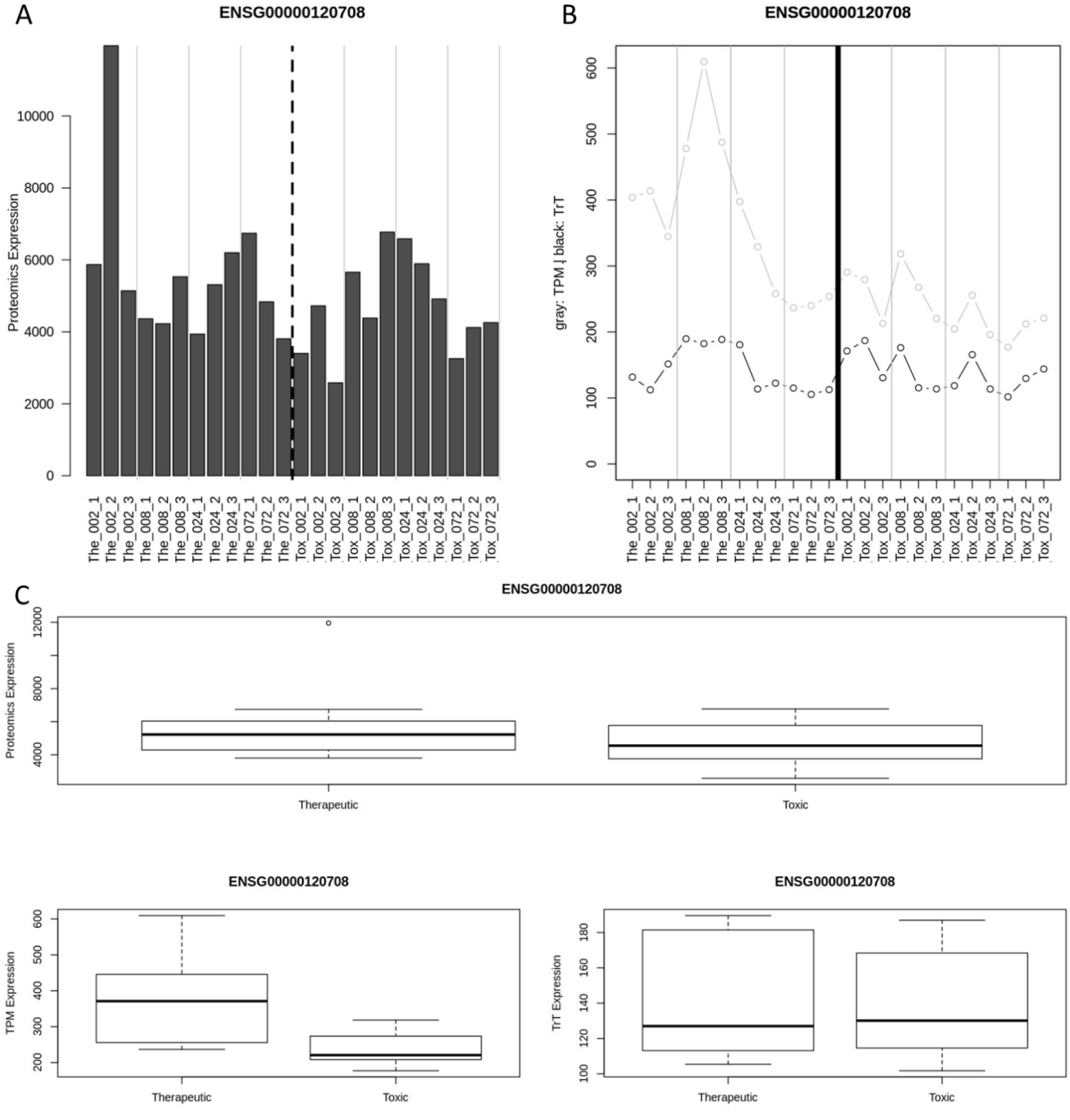
Expression values of TGFBI. **A:** Proteomics expression values. **B:** TPM and TrT expression values: Gray: TPM, Black: TrT. **C:** Boxplots of the expression values for Proteomics, TPM, and TrT. TPM: Transcripts Per Million. TrT: Translatable Transcripts.

### Biological Interpretation (GOrilla)

After observing the effects TrT could have, we investigated whether those could also affect the biological interpretation. Specifically, we focused on comparing the enriched gene ontology-sets (GO-sets) of both TPM and TrT via GOrilla. The results showed that the number of GO terms was always higher in TPM than TrT (Figure 6). Such a decrease in GO terms for TrT, though, was not related to a lower number of DEGs from such quantifier. When analyzing the commonalities and differences between TPM and TrT across compounds, we observed 2 different behaviors (Figure 6). For some compounds (5FU, AMI, EPI & MXT), the number of TrT GO terms shared with TPM was either greater or equal than the exclusive ones, while most of the TPM terms were exclusive. On the other compounds, the GO terms shared between them were rather the minority for both sets.

**Figure 6:**
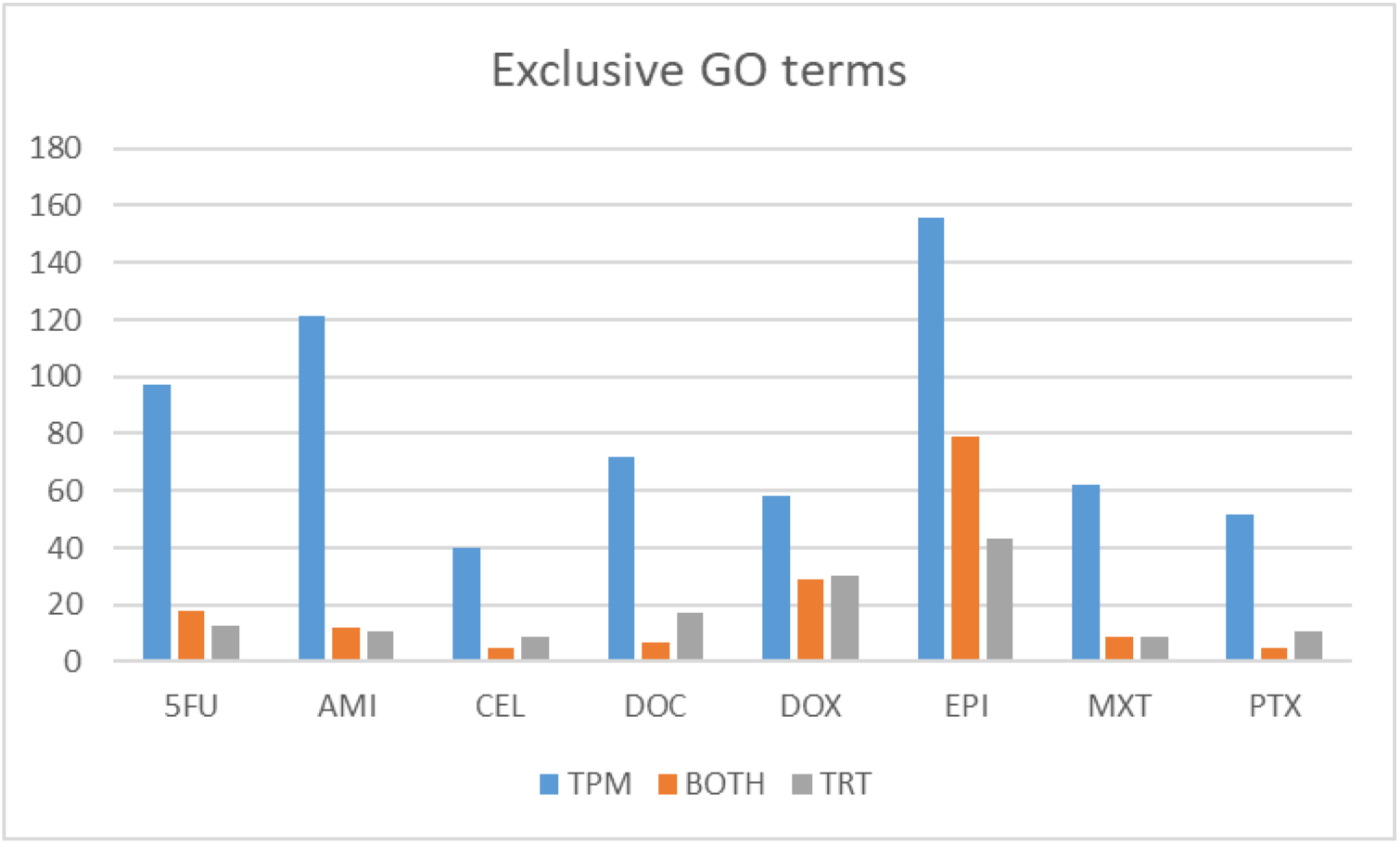
Number of exclusive GO terms for each quantifier (TPM & TRT) and the ones included in both (BOTH). TPM: Transcripts Per Million. TrT: Translatable Transcripts. 5FU: Fluorouracil. AMI: Amiodarone. CEL: Celecoxib. DOC: Docetaxel. DOX: Doxorubicin. EPI: Epirubicin. MXT: Mitoxantrone. PTX: Paclitaxel.An example of how the misclassification of a DEG may affect the biological interpretation of a comparison (with enough genes being misclassified) was the Cardiac-Specific Homeo Box (NKX2-5) gene. According to TPM, expression was significantly higher in the UNTR samples when compared to its corresponding samples in the therapeutic dose in 5FU, AMI, and DOC, and part of the Cardiac Muscle Tissue Morphogenesis enriched GO set of genes. TrT, though, showed no difference between both scenarios (Figure 7A). This meant that the miRNA regulation was predicted to be stronger in the control samples than the treatment ones, to the point where the difference between both conditions was not significant. To contrast that prediction, we searched for all 18 MTIs with either weak or strong evidence of their inhibitory regulation. We observed that the expression of all those miRNAs in the Control samples was greater or equal than the therapeutic ones, confirming the TrT prediction (Figure 7B).

**Figure 7:**
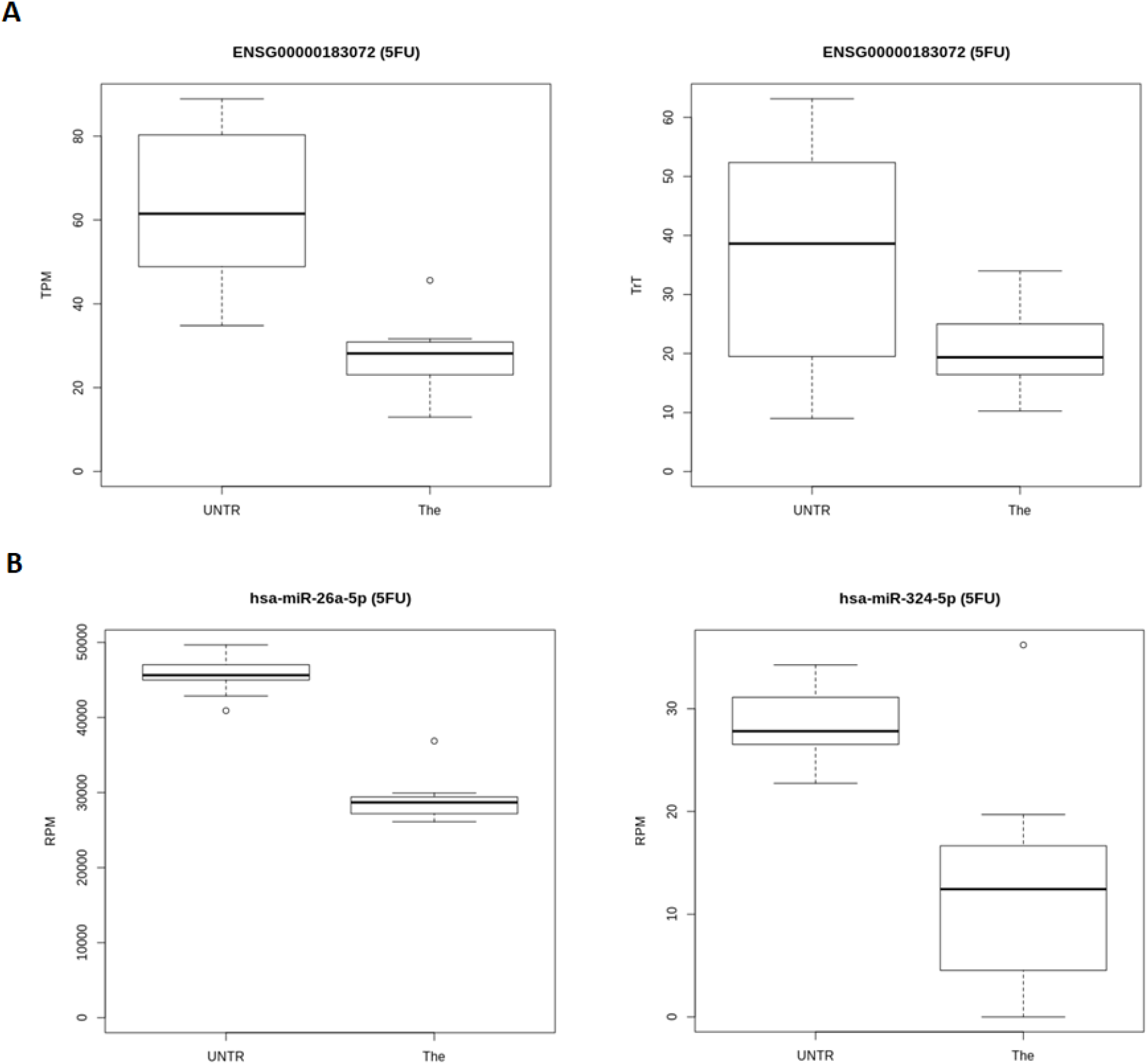
NKX2-5 gene: Expression and Regulation between Untreated (UNTR) and Therapeutic (The) samples treated with Fluorouracil (5FU). **A**. TPM (left) and TrT (right) expression represented in boxplots. **B**. Examples of the expression in Reads Per Million (RPM) of miRNAs that regulate the NKX2-5 gene.

### Sensitivity, Specificity, and Accuracy results in TPM and TRT

So far, we saw that, at least in some cases, TrT could be a better proxy for proteomics than TPM, and that the biological interpretation was simplified when TrT was applied. Afterward, we wanted to assess the global accuracy of TrT and TPM by comparing it to proteomics.

We calculated the accuracy of both transcriptomics values for all compounds in 3 different comparisons: therapeutic versus toxic doses, untreated control versus therapeutic dose, and untreated control versus toxic dose. To evaluate the differential expression of the last 2 comparisons, we used the same methodology as the one used for therapeutic versus toxic doses.

We observed an ambiguous effect of TrT in relation to TPM in all 3 comparisons from a global perspective (Figure 8). Individually, though, we observed a compound effect reflected on different accuracy differences for each of the 8 compounds, which could be categorized into 3 groups when analyzing the accuracy differences for each compound. Most of them showed ambiguous results, in which both improvement and decline of the accuracy happened. EPI, instead, showed a continuous improvement across all comparisons. The last group, contrarily (CEL & MXT), showed a continuous decline across all comparisons.

**Figure 8:**
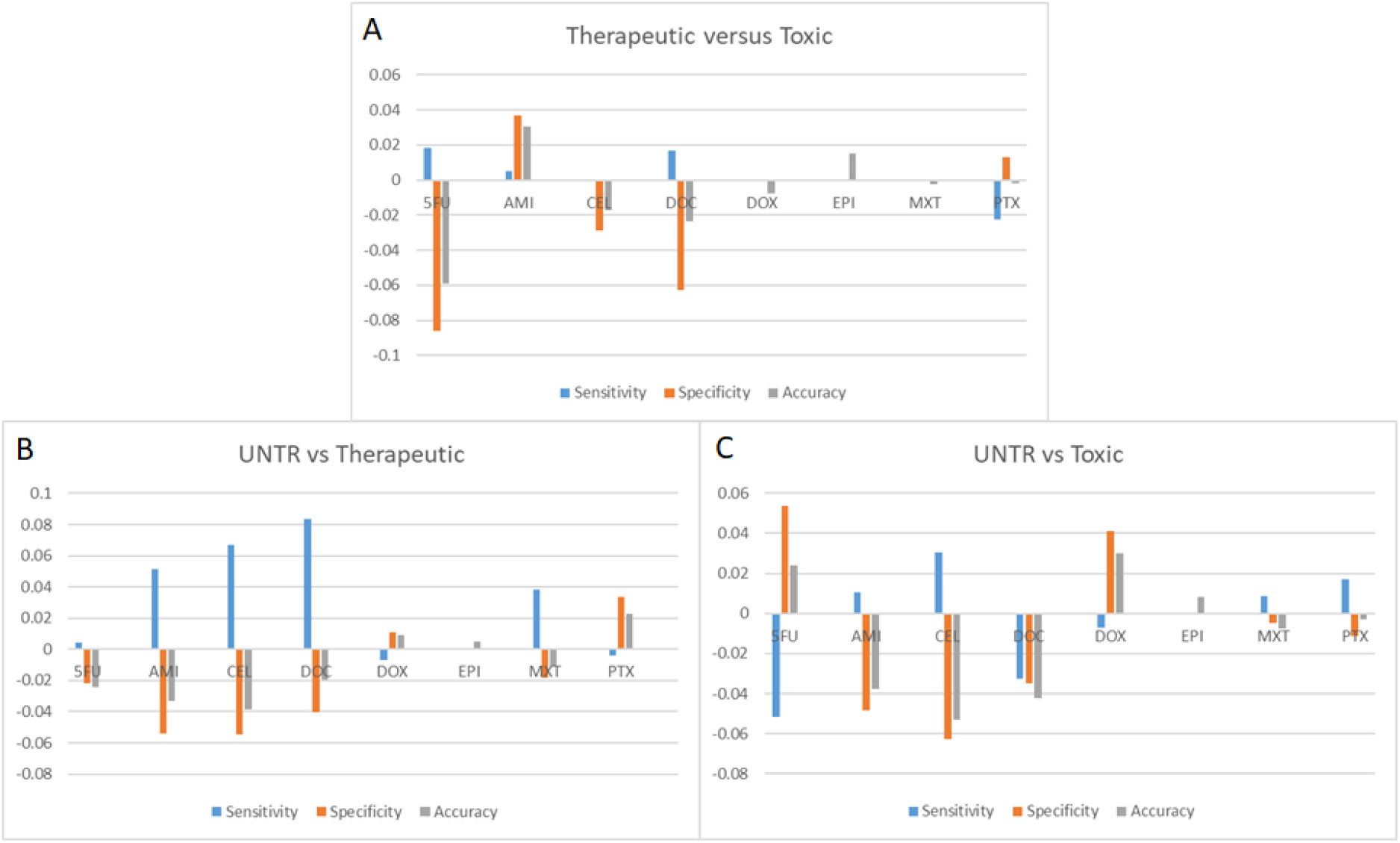
Difference in Sensitivity, Specificity, and Accuracy when changing the quantifier from TPM to TrT, each comparison encompassing 8 compounds: Fluorouracil (5FU), Amiodarone (AMI), Celecoxib (CEL), Docetaxel (DOC), Doxorubicin (DOX), Epirubicin (EPI), Mitoxantrone (MXT), and Paclitaxel (PTX). A: Therapeutic versus Toxic doses. B: Untreated versus Therapeutic dose. C: Untreated versus Toxic dose.

## Discussion

Correlation between transcriptomics and proteomics has always tended to be low. This is true both in previous and in the current study, proving transcriptomics as a poor proxy of the phenotypical changes of the cell. Trying to predict the regulatory effect of miRNA (TrT) to improve such proxy (TPM) proved beneficial for a portion of the gene set. This is unsurprising, knowing that not all genes are regulated by miRNAs, and only a small portion of proteins are quantified through the proteomics pipeline. TrT also showed a simplified biological interpretation of the changes across conditions, although its results compared to TPM varied compound-wise.

We did not only take miRNAs into account, though. Other ncRNAs, such as circRNAs (although recent studies have observed some of them as coding), were also quantified. Due to the novelty of circRNAs, in contrast with other Omics, there was no golden standard database to refer to when seeking their specific IDs nor the way they should be quantified, among other reasons, because the existing databases were seldom updated^18,49^ nor online^50^. One of the most recent recommendations^44^ in these circumstances was the use of at least 2 different circRNA identification software tools. The reason behind this procedure was based on the flawed accuracy of these programs due to their inherent biases. Fortunately, even if this bias differed for every tool, it had been shown that the convergence between 2 tools offered a much lower proportion of false positives in contrast to their original population of predictions^44^.

We decided to pool all samples so that we had the maximum sequencing depth available for the identification process. Otherwise, selecting a random sample would lead to an under-representation of some circRNA molecules, which would not be identified as such, even if they would be in the rest of the samples.

The argumentation behind the selection of CIRI2 and circExplorer2 was twofold. First, their accuracy outcomes in previous reviews portrayed them to be some of the best predictors currently available. Secondly, our comparison of several predictors revealed that both output a substantial proportion of molecules that were also outputted by other predictors (low false-positive rate), without sacrificing sensitivity. This was not the case for circRNA_finder^32^, which found several times more molecules than the other predictors with a very low overlap (∼7%). find_circ^18^, although having a similar proportion of overlap as circExplorer2 and CIRI2, did not present such a good accuracy in previous studies.

The model we applied considered several factors in the post-transcriptional regulation. We built it by trying to reach an equilibrium at the complexity level: enough complexity to maximize the representation of the molecular reality of the cell; but without including so many variables that would have made the model either unbuildable or unfeasible to run in practical terms. Our model also reflected how little is still known about the post-transcriptional regulation, and recent discoveries are constantly being made public around it^42,51^.

This is especially the case with miRNA regulation, where the complementarity and regulatory strength seems to not only be dependent on the seed region but other loci such as the 3’ end of miRNAs^42^. These new observations are currently being incorporated into the interaction predictive tools, making probably in the future the more classical tools obsolete.

The genes exemplified as good proxies for proteomics are indeed regulated by miRNAs. MYH9 is a non-muscle myosin chain IIA gene which plays a major role in early mammalian development, while in the adult heart is only expressed in the non-myocyte cells^52^. Several miRNAs have been either positively or negatively associated with the gene’s expression^53^. For TGFBI, miRNA regulation has also been shown, to the point of even being related to chemoresistance^54^.

Our model was based on a miRNA regulation on a 1-on-1 basis: if a miRNA can bind a transcript (after taking into account the endogenous competition), the latter will not be able to be translated. Other recent models prefer to focus on the targeting efficacy (as a function of the affinity between the two molecules) while ignoring the ceRNAs^51^. We acknowledge that both models lack each other’s strengths, and a combination of the two would have been optimal.

In addition to that, there could have been other important factors that could have been taken into account. One example could be the translation ratio, i.e., the number of proteins translated per transcript, which depends on the number of ribosomes that may bind to it, and its half-life. Indeed, half-lives, both of the transcript and the protein, makes it difficult to transient from one OMICs to another: the increase in proteomics might be by an increase in transcriptomics with a short half-life that cannot be seen, or a long half-life of proteins may delay the decrease in their abundance due to a decrease in translational output.

Proteomics, although being the technology used as a reference to represent the phenotypical changes in the cell, is quite limited in achieving so. This is especially the case due to its lack of sensitivity, where generally only around 1000-1500 proteins are quantified, making the analysis of the proteome quite challenging. In addition to that, the output scale of such quantifications can be quite dramatic when the technology or machinery involved changes.

Our next steps would include trying to breach through the gap between both OMICs by, not only including important parameters that can improve the proteomics prediction, but also changing the algorithms involved in the model, namely, using a supervised quantitative machine learning model. That way, the use of new parameters is easier to both implement and evaluate.This is specially relevant for the imputation of proteomics data, prone to missing values due to the stochastic nature of mass spectrometry. Our work concerning this strategy can be found in our published article^57^.

We created a formula to model the post-transcriptional regulation and its effects on protein expression. We assessed its behavior in both general and specific cases. As expected, due to the variability of each gene’s regulation, the formula benefitted only a subset of genes. We portrayed the importance of reaching a more accurate description of the cellular changes, and how far we still are from that objective. Therefore, we must not simply continue with the most popular technology, but try to reach a better approach out of the current era of big data.

## Supporting information

Supplementary Table GO terms 2

Supplementary Table GO terms 1

## Bibliography

1. Zhang, Z., Wu, S., Stenoien, D. L. & Paša-Tolić, L. High-throughput proteomics. Annu. Rev. Anal. Chem. (Palo Alto. Calif). 7, 427–54 (2014).

2. Peng, J., Elias, J. E., Thoreen, C. C., Licklider, L. J. & Gygi, S. P. Evaluation of multidimensional chromatography coupled with tandem mass spectrometry (LC/LC-MS/MS) for large-scale protein analysis: The yeast proteome. J. Proteome Res. 2, 43–50 (2003).

3. Washburn, M. P., Wolters, D. & Yates, J. R. Large-scale analysis of the yeast proteome by multidimensional protein identification technology. Nat. Biotechnol. 19, 242–247 (2001).

4. Cagney, G. et al. Human tissue profiling with multidimensional protein identification technology. J. Proteome Res. 4, 1757–67 (2005).

5. Chen, G. et al. Discordant protein and mRNA expression in lung adenocarcinomas. Mol. Cell. Proteomics 1, 304–13 (2002).

6. Lemée, J.-M. et al. Integration of transcriptome and proteome profiles in glioblastoma: looking for the missing link. BMC Mol. Biol. 19, 13 (2018).

7. Rogers, S. et al. Investigating the correspondence between transcriptomic and proteomic expression profiles using coupled cluster models. Bioinformatics 24, 2894–2900 (2008).

8. Dhingra, V., Gupta, M., Andacht, T. & Fu, Z. F. New frontiers in proteomics research: A perspective. International Journal of Pharmaceutics vol. 299 1–18 (2005).

9. Belle, A., Tanay, A., Bitincka, L., Shamir, R. & O’Shea, E. K. Quantification of protein half-lives in the budding yeast proteome. Proc. Natl. Acad. Sci. U. S. A. 103, 13004–13009 (2006).

10. Baek, D. et al. The impact of microRNAs on protein output. Nature 455, 64–71 (2008).

11. Selbach, M. et al. Widespread changes in protein synthesis induced by microRNAs. Nature 455, 58–63 (2008).

12. Ambros, V. The functions of animal microRNAs. Nature 431, 350–355 (2004).

13. Lewis, B. P., Shih, I., Jones-Rhoades, M. W., Bartel, D. P. & Burge, C. B. Prediction of Mammalian MicroRNA Targets. Cell 115, 787–798 (2003).

14. Lim, L. P. et al. Microarray analysis shows that some microRNAs downregulate large numbers of-target mRNAs. Nature 433, 769–773 (2005).

15. Zaphiropoulos, P. G. Exon skipping and circular RNA formation in transcripts of the human cytochrome P-450 2C18 gene in epidermis and of the rat androgen binding protein gene in testis. Mol. Cell. Biol. 17, 2985–2993 (1997).

16. Chen, L. L. & Yang, L. Regulation of circRNA biogenesis. RNA Biol. 12, 381–388 (2015).

17. Jeck, W. R. et al. Circular RNAs are abundant, conserved, and associated with ALU repeats. RNA 19, 141–157 (2013).

18. Memczak, S. et al. Circular RNAs are a large class of animal RNAs with regulatory potency. Nature 495, 333–338 (2013).

19. Rong, D. et al. An emerging function of circRNA-miRNAs-mRNA axis in human diseases. Oncotarget 8, (2017).

20. Wang, K. et al. The long noncoding RNA CHRF regulates cardiac hypertrophy by targeting miR-489. Circ. Res. 114, 1377–88 (2014).

21. Salmena, L., Poliseno, L., Tay, Y., Kats, L. & Pandolfi, P. P. A ceRNA hypothesis: the Rosetta Stone of a hidden RNA language? Cell 146, 353–8 (2011).

22. Kuepfer, L. et al. A model-based assay design to reproduce in vivo patterns of acute drug-induced toxicity. Archives of Toxicology vol. 92 553–555 (2018).

23. Bolger, A. M., Lohse, M. & Usadel, B. Trimmomatic: a flexible trimmer for Illumina sequence data. Bioinformatics 30, 2114–20 (2014).

24. Baras, A. S. et al. miRge - A Multiplexed Method of Processing Small RNA-Seq Data to Determine MicroRNA Entropy. PLoS One 10, e0143066 (2015).

25. Langmead, B., Trapnell, C., Pop, M. & Salzberg, S. L. Ultrafast and memory-efficient alignment of short DNA sequences to the human genome. Genome Biol. 10, R25 (2009).

26. Zhang, X.-O. et al. Diverse alternative back-splicing and alternative splicing landscape of circular RNAs. Genome Res. 26, 1277–87 (2016).

27. Gao, Y., Zhang, J. & Zhao, F. Circular RNA identification based on multiple seed matching. Brief. Bioinform. 19, 803–810 (2018).

28. Li, H. & Durbin, R. Fast and accurate short read alignment with Burrows-Wheeler transform. Bioinformatics vol. 25 1754–1760 https://pubmed.ncbi.nlm.nih.gov/19451168/ (2009).

29. Quinlan, A. R. & Hall, I. M. BEDTools: a flexible suite of utilities for comparing genomic features. Bioinformatics 26, 841–2 (2010).

30. ENSEMBL. FTP Download. https://www.ensembl.org/info/data/ftp/index.html (2020).

31. Patro, R., Duggal, G., Love, M. I., Irizarry, R. A. & Kingsford, C. Salmon provides fast and bias-aware quantification of transcript expression. Nat. Methods 14, 417–419 (2017).

32. Westholm, J. O. et al. Genome-wide Analysis of Drosophila Circular RNAs Reveals Their Structural and Sequence Properties and Age-Dependent Neural Accumulation. Cell Rep. 9, 1966–1980 (2014).

33. Griffiths-Jones, S., Grocock, R. J., van Dongen, S., Bateman, A. & Enright, A. J. miRBase: microRNA sequences, targets and gene nomenclature. Nucleic Acids Res. 34, D140–D144 (2006).

34. Durinck, S. M. Y. et al. BioMart and Bioconductor: a powerful link between biological databases and microarray data analysis. Bioinformatics 21, 3439–3440 (2005).

35. Li, H. et al. The Sequence Alignment/Map format and SAMtools. Bioinformatics 25, 2078–2079 (2009).

36. John, B. et al. Human microRNA targets. PLoS Biol. 2, (2004).

37. R Core Team. R: A Language and Environment for Statistical Computing. (2019).

38. RStudio Team. RStudio: Integrated Development Environment for R. (2019).

39. Wei, T. & Simko, V. R package ‘corrplot’: Visualization of a Correlation Matrix. (2017).

40. Analyzing RNA-seq data with DESeq2. https://bioconductor.org/packages/release/bioc/vignettes/DESeq2/inst/doc/DESeq2.html#why-un-normalized-counts.

41. Robinson, M. D. & Oshlack, A. A scaling normalization method for differential expression analysis of RNA-seq data. Genome Biol. 11, 1–9 (2010).

42. Chipman, L. B. & Pasquinelli, A. E. miRNA Targeting: Growing beyond the Seed. Trends in Genetics vol. 35 215–222 (2019).

43. Chou, C. H. et al. MiRTarBase update 2018: A resource for experimentally validated microRNA–target interactions. Nucleic Acids Res. (2018) doi:10.1093/nar/gkx1067.

44. Hansen, T. B. Improved circRNA identification by combining prediction algorithms. Front. Cell Dev. Biol. 6, 20 (2018).

45. Bartel, D. P. MicroRNAs: target recognition and regulatory functions. Cell 136, 215–33 (2009).

46. Arvey, A., Larsson, E., Sander, C., Leslie, C. S. & Marks, D. S. Target mRNA abundance dilutes microRNA and siRNA activity. Mol. Syst. Biol. 6, 363 (2010).

47. Franco-Zorrilla, J. M. et al. Target mimicry provides a new mechanism for regulation of microRNA activity. Nat. Genet. 39, 1033–7 (2007).

48. MS, E., JR, N. & PA, S. MicroRNA Sponges: Competitive Inhibitors of Small RNAs in Mammalian Cells. Nat. Methods 4, (2007).

49. Li, J. H., Liu, S., Zhou, H., Qu, L. H. & Yang, J. H. StarBase v2.0: Decoding miRNA-ceRNA, miRNA-ncRNA and protein-RNA interaction networks from large-scale CLIP-Seq data. Nucleic Acids Res. 42, D92–7 (2014).

50. Liu, Y. C. et al. CircNet: A database of circular RNAs derived from transcriptome sequencing data. Nucleic Acids Res. 44, D209–D215 (2016).

51. McGeary, S. E. et al. The biochemical basis of microRNA targeting efficacy. Science (80-.). 366, eaav1741 (2019).

52. Ma, X. & Adelstein, R. S. In vivo studies on nonmuscle myosin II expression and function in heart development. Front. Biosci. 17, 545–555 (2012).

53. Yu, M. et al. Prognostic impact of MYH9 expression on patients with acute myeloid leukemia. Oncotarget 8, 156–163 (2017).

54. Bissey, P. A. et al. Dysregulation of the MiR-449b target TGFBI alters the TGFβ pathway to induce cisplatin resistance in nasopharyngeal carcinoma. Oncogenesis 7, (2018).

55. Nakamoto, M., Jin, P., O’Donnell, W. T. & Warren, S. T. Physiological identification of human transcripts translationally regulated by a specific microRNA. Hum. Mol. Genet. 14, 3813–3821 (2005).

56. Moore, M. J. et al. MiRNA-target chimeras reveal miRNA 3′-end pairing as a major determinant of Argonaute target specificity. Nat. Commun. 6, 1–17 (2015).

57. Ochoteco Asensio, J., Verheijen, M. & Caiment, F. Predicting missing proteomics values using machine learning: Filling the gap using transcriptomics and other biological features. Comput Struct Biotechnol J 20, 2057–2069, doi:10.1016/j.csbj.2022.04.017 (2022).

